# OT-Fusion: Cortex-conditioned Fusion of Matched PET and MRI Images Using Optimal Transport

**DOI:** 10.1101/2025.03.02.641024

**Authors:** Emir Türkölmez, Yasir Ali Khan, Caroline Bund, Izzie Jacques Namer, A. Ercüment Çiçek, Selim Aksoy

**Affiliations:** Department of Computer Engineering, Bilkent University, Ankara, Türkiye; Department of Nuclear Medicine and Molecular Imaging, Institut de Cancérologie Strasbourg Europe (ICANS), Strasbourg, France

**Author notes:** Correspondence: {, }. These authors have contributed equally (Emir Türkölmez, Yasir Ali Khan).

**Keywords:** PET, MRI, Image fusion, Optimal transport

## Abstract

Positron Emission Tomography (PET) and Magnetic Resonance Imaging (MRI) are crucial in diagnosing various medical conditions related to brain. FDG-PET provides essential functional information by capturing synaptic transmission activity, while MRI offers anatomical details. The integration of these modalities has become increasingly valuable, especially with the advent of hybrid PET/MRI systems that enable acquisition of both images in a single session. However, a notable challenge arises due to the inherent differences in spatial resolution between PET and MRI. PET images often have lower spatial resolution, leading to metabolic signals from active regions, such as the cerebral cortex and basal ganglia, appearing diffused and extended into adjacent areas like the white matter. This diffusion can complicate the accurate localization of metabolic activity, posing difficulties in clinical assessments, particularly in neurological pathologies such as epilepsy where precise mapping is crucial. To address this issue, we propose a novel approach that effectively maps the metabolic activity observed in PET onto the cortical structures delineated in MRI. We formulate this mapping as an optimal transport problem that corresponds to finding the optimum way to transform one probability distribution into another. Our method aims to enhance the structural quality of PET images, making them more detailed and informative for clinical decisions related to neurological disorders. The experiments using paired PET and MRI images indicate that our method performs better than the state-of-the-art image fusion methods on quantitative metrics and improves image quality based on visual qualitative evaluation by an expert physician.

## 1 Introduction

Positron emission tomography (PET) and magnetic resonance imaging (MRI) are two medical imaging modalities that offer complementary information. While PET captures metabolic activity, MRI provides anatomical details. Recently, simultaneous PET/MRI has gained popularity, enabling the acquisition of both images in a single scan. However, PET often has lower spatial resolution than MRI, meaning a voxel in PET corresponds to multiple voxels in the matched MRI image. This discrepancy poses a challenge: Activity recorded in a PET voxel represents an average over multiple structures visible in the MRI, leading to a diffused PET image and complicating the identification of active regions.

This issue is particularly significant in brain imaging, where precise evaluation is essential. In a matched PET/MRI brain scan, metabolic and/or synaptic transmission activity captured by PET with FDG radiotracer is expected to be observed in the cortical regions and basal ganglia. However, due to the lower resolution of PET, this activity often appears to extend beyond the cortex into the white matter, which should be inactive. This is critical in conditions like epilepsy, where accurate localization of hypometabolic regions is necessary to determine epileptiform zone susceptible to surgery. Therefore, a method is needed to map the activity observed in the dispersed PET image onto the cortex delineated in the high-resolution MRI.

Related approaches in the literature may not address this challenge. Image super-resolution models have been used to increase the resolution of PET images [7]. However, training such models requires high-resolution versions of PET images that are rarely available, if at all. As an alternative, image fusion techniques map PET activity on the MRI images [8,9]. Nevertheless, they do not improve the resolution of PET but simply blend it with MRI. They also cannot distinguish cortical regions. Therefore, there is still a need for a method that fuses the contributions from both data sources while enhancing the resolution of PET for clinical assessment.

In this paper, we propose a novel method, OT-Fusion, for mapping the metabolic activity captured on a brain PET scan onto the cortical regions in the corresponding MRI image. OT-Fusion does not require high-resolution versions of the PET image and moves the total activity captured on PET to the cortex on MRI in line with the underlying biology. We formulate this mapping as an *optimal transport* problem that corresponds to the process of finding the most efficient way to transform one probability distribution (PET activity) into another (cortex in the MRI image). Using a dataset of matched PET and MRI images of healthy individuals and patients with conditions such as Hashimoto’s encephalopathy, Alzheimer’s and Epilepsy, we show that OT-Fusion results in higher quality images compared to state-of-the-art fusion methods with respect to quantitative image quality metrics. We also present qualitative results where OT-Fusion is able to capture the low-activity cortex regions more accurately than the state-of-the-art which predict high activity over white matter and regions affected by disease. The code and data will be released upon acceptance of the manuscript.

## 2 Related Work

Image Super-Resolution (SR) aims to enhance the resolution of an image by reconstructing a high-resolution (HR) version from a low-resolution (LR) input. For instance, Song et al. [7] adapt a shallow architecture from Dong et al. [2] and a very deep architecture from Kim et al. [4] to develop a PET SR model. They incorporate HR MRI images and auxiliary information such as radial and axial locations of pixels. Ozaltan et al. [5] also use a model with convolutional blocks with residual connections for efficient training. The main bottleneck is that they require an HR target image during training which is rarely available, if at all.

Image Fusion aims to superpose and blend two images to produce a single enhanced version with information from both modalities. This method does not require an HR version of PET unlike SR. Xu et al. [9] propose an end-to-end network named U2Fusion that estimates the relative contribution of each image and adaptively preserves the similarity between the fusion result and the source images. The fusion of PET and MRI data is obtained by enhancing the overall content with predefined fusion rules. Later, Xu and Ma [8] develop a multi-modal framework named EMFusion that uses a model for transforming input images into a common feature space, and another model that combines these features using both surface-level (e.g., salience and abundance) and deep-level (e.g., feature uniqueness) constraints. Unfortunately, such models only learn to stress and enhance visibility of certain features of one modality in specific regions of interest in the target, while downplaying others. They do not improve resolution or align the original activity distribution in PET with the target MRI image.

## 3 Methodology

### 3.1 Dataset

We use simultaneously acquired PET/MRI images of 14 healthy individuals (controls) and 5 individuals (patients) with Hashimoto’s encephalopathy (1), Alzheimer’s (1) and Epilepsy (3). The images are taken using a General Electric Healthcare Signa PET/MRI scanner with 18F-FDG injection at the Hautepierre Hospital — University Hospitals of Strasbourg. Individuals underwent imaging 30 minutes after the injection of 2 MBq/kg 18F-FDG, in accordance with current EANM recommendations. Prior to injection, individuals fasted for 6 hours, and blood glucose levels (< 1.3 g/L) were verified using a glucometer. Total PET/MRI acquisition time was 15 minutes.

Time-of-flight (TOF) PET images were acquired in 3D mode with a 384 × 384 pixel matrix and a pixel size of 2.7 mm, using iterative ordered subset expectation maximization (OSEM) reconstruction. All MRI images were also acquired in 3D mode (T1 gradient echo, 3D T2, 3D fluid-attenuated inversion recovery (FLAIR), arterial spin labeling (ASL) sequences) with an isotropic resolution of 1 mm. There are 744 slice pairs (53 slice pairs/person on average) in the control and 327 slice pairs (65 slice pairs/person on average) in the patient data.

The data for the healthy individuals are obtained as part of the clinical trial NCT05355857^3^. All participants provided written informed consent. The Ethics Committee approved the ATLATEP study in (no. SI 22.00287.000073, March 22, 2022). Informed consent for using clinico-radiological data for research purposes was also obtained from the patients. Institutional review board approval was obtained for the INTARTEP study and registered under IRB-2025-2.

### 3.2 Preprocessing

We crop the centered 300 × 300 pixel regions from the MRI slices. We also crop the corresponding region from each PET slice and use interpolation with the Lanczos kernel to increase its resolution to the MRI level. Each slice pair in the matched PET/MRI images is processed independently in 2D.

We use the DL+DiReCT framework of Rebsamen et al. [6], which combines deep learning-based neuroanatomy segmentation and cortex parcellation with diffeomorphic registration, to measure cortical thickness from T1-weighted MRI images. This method segments the brain’s anatomical structures using a CNN and differentiates cortical regions. This step (*cortex prediction*) produces a segmentation map where each pixel is associated with a probability of belonging to the cortical region in the MRI image (higher probability = brighter pixel intensity). Then, we employ morphological operations to obtain a mask of the brain region from the PET image and use this mask to remove the skull regions from all inputs including the cortex prediction. Finally, we apply a threshold, empirically selected as 0.4, to the cortex prediction image to obtain a binary cortex mask.

### 3.3 Optimal Transport Formulation

We aim to map the functional activity (represented by the intensity distribution in the PET space) onto the cortex regions (represented by the structural prediction in the MRI space). We formulate this mapping as an optimal transport (OT) problem that corresponds to the process of finding the most efficient way to transform one probability distribution into another.

Let *X* = {*x*_1_, …, *x*_*m*_} and *Y* = {*y*_1_, …, *y*_*n*_} be the sets of pixels in the matched PET and MRI images, respectively. We define function 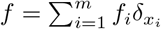 where 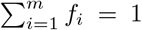 and *f*_*i*_ ≥ 0 as the discrete probability distribution that represents the metabolic activity at each pixel of the PET image. *δ*_*x*_ denotes the where and Dirac delta function. Similarly, the function 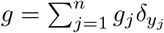where 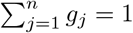 and *g*_*j*_ ≥ 0 represents the presence of the cortex at each pixel of the MRI image. These functions are obtained by normalizing the intensity values in the PET image and the cortex prediction in the MRI image, respectively. The goal is to find the minimal total cost solution of the transport problem where the cost of moving a unit mass from pixel *x*_*i*_ to pixel *y*_*j*_ is defined as a function of the distance traveled as *C*(*x*_*i*_, *y*_*j*_) = ∥*x*_*i*_ − *y*_*j*_∥ _2_. While alternative distance definitions are possible, we use the Euclidean distance between pixel pairs in this paper.

The OT problem seeks to minimize the transport cost over all possible movement patterns using the following formulation:

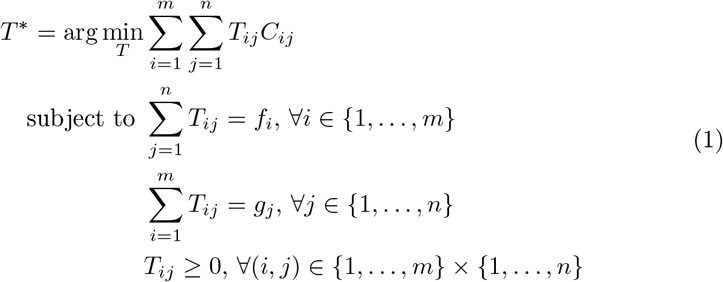

where *C*_*ij*_ = *C*(*x*_*i*_, *y*_*j*_) and 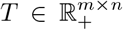 with *T*_*ij*_ = *T* (*x*_*i*_, *y*_*j*_) is the transport plan that corresponds to the amount of mass to be moved between *x*_*i*_ and *y*_*j*_. The constraints ensure that the marginal distributions of the transport plan match the distributions *f* and *g*. In this plan, each pixel in the PET image can distribute its activity onto multiple cortex pixels in MRI, and each pixel in the cortex prediction can receive activities from multiple pixels in PET.

We use the network simplex method that models the transport as flow through a bipartite graph that connects source nodes to target nodes, as described in [1], for the solution of the OT problem. We use the polynomial-time implementation of this method in the Python Optimal Transport library [3].

To add further flexibility to the solution, we introduce a parameter *α* that forces a particular percentage of the mass at each pixel not to be moved. First, both the source and target distributions are scaled as *f* ^*′*^ = (1 − *α*)*f* and *g*^*′*^ = (1 − *α*)*g*. Then, the solution to the OT problem is obtained as *T* ^*′*^ by using *f* ^*′*^ and *g*^*′*^ in the formulation in (1). Finally, the transport plan is computed as *T* = *T* ^*′*^ +diag(*αf*) where the “diag” operation creates a matrix with the elements of the input vector along the diagonal. This assumes that the source and target images *f* and *g*, respectively, have the same number of pixels and are aligned as in our case. As two extreme scenarios, when *α* = 0, all mass (metabolic activity) is transported from the source to the target, and when *α* = 1, no transportation is performed. This force parameter *α* aims to control the rate of the transport, and can potentially account for the uncertainty in the cortex prediction step from the MRI data. In all cases, the total metabolic activity in the input PET image is preserved after the mapping.

## 4 Experiments

### 4.1 Quantitative Evaluation

We measure the performance of the proposed methodology by using peak signal-to-noise ratio (PSNR), structural similarity index (SSIM), and learned perceptual image patch similarity (LPIPS). The input PET images are used as the reference in the computation of these metrics as also done by Xu et al. [9] and Xu and Ma [8]. The *α* parameter is empirically set as 0.65 in the analyses presented in this section.

Table 1 shows the quantitative results for OT-Fusion, U2Fusion, and EMFusion using 744 slice pairs from the control and 327 slice pairs from the patient data. We train the U2Fusion model with our control data using the code in the corresponding repository^4^ and use the pretrained EMFusion model obtained from the corresponding repository^5^ in these experiments. The OT-Fusion output has a higher PSNR (higher is better) than both U2Fusion and EMFusion, indicating better signal fidelity relative to the PET images. Similarly, SSIM is higher (higher is better) for OT-Fusion compared to both U2Fusion and EMFusion, reflecting its closer resemblance to the original PET data. The LPIPS value for OT-Fusion is lower (lower is better) than both U2Fusion and EMFusion, indicating that OT-Fusion preserves perceptual similarity to the target images more effectively, resulting in a more visually coherent and structurally accurate reconstruction.

**Table 1.** Quantitative image quality metrics (average and standard deviation) computed when the original PET image is used as the reference. Best results are in bold.

| Method | Healthy (control) individuals |  |  | Patients |  |  |
| --- | --- | --- | --- | --- | --- | --- |
| | PSNR (↑) | SSIM (↑) | LPIPS (↓) | PSNR (↑) | SSIM (↑) | LPIPS (↓) |
| **OT-Fusion** | **21.18 ± 3.36** | **0.82 ± 0.09** | **0.14 ± 0.06** | **21.42 ± 4.19** | **0.81 ± 0.09** | **0.18 ± 0.05** |
| U2Fusion[9] | 17.02 ± 1.78 | 0.74 ± 0.08 | 0.20 ± 0.05 | 18.04 ± 4.02 | 0.73 ± 0.12 | 0.25 ± 0.07 |
| EMFusion[8] | 15.20 ± 1.75 | 0.73 ± 0.08 | 0.20 ± 0.06 | 16.92 ± 4.37 | 0.71 ± 0.13 | 0.26 ± 0.07 |

### 4.2 Qualitative Evaluation

For qualitative evaluation, we visually compared axial slices of FDG PET images (using both B&W and French LUT color map), T1-weighted MRI, MRI cortex, binary cortex, U2Fusion, EMFusion, and OT-Fusion images for six individuals (Figure 1). The first three columns represent healthy volunteers, while the last three columns correspond to patients with distinct pathologies: Hashimoto’s encephalopathy, Alzheimer’s dementia, and cortical dysplasia leading to epileptic seizures. All pathological cases exhibit bilateral (columns 4 and 5) or unilateral (column 6) parietal cortical hypometabolism on the original PET scans. Highlighted regions in each case are enclosed within red bounding boxes and enlarged in Figure 2.

**Fig. 1.**
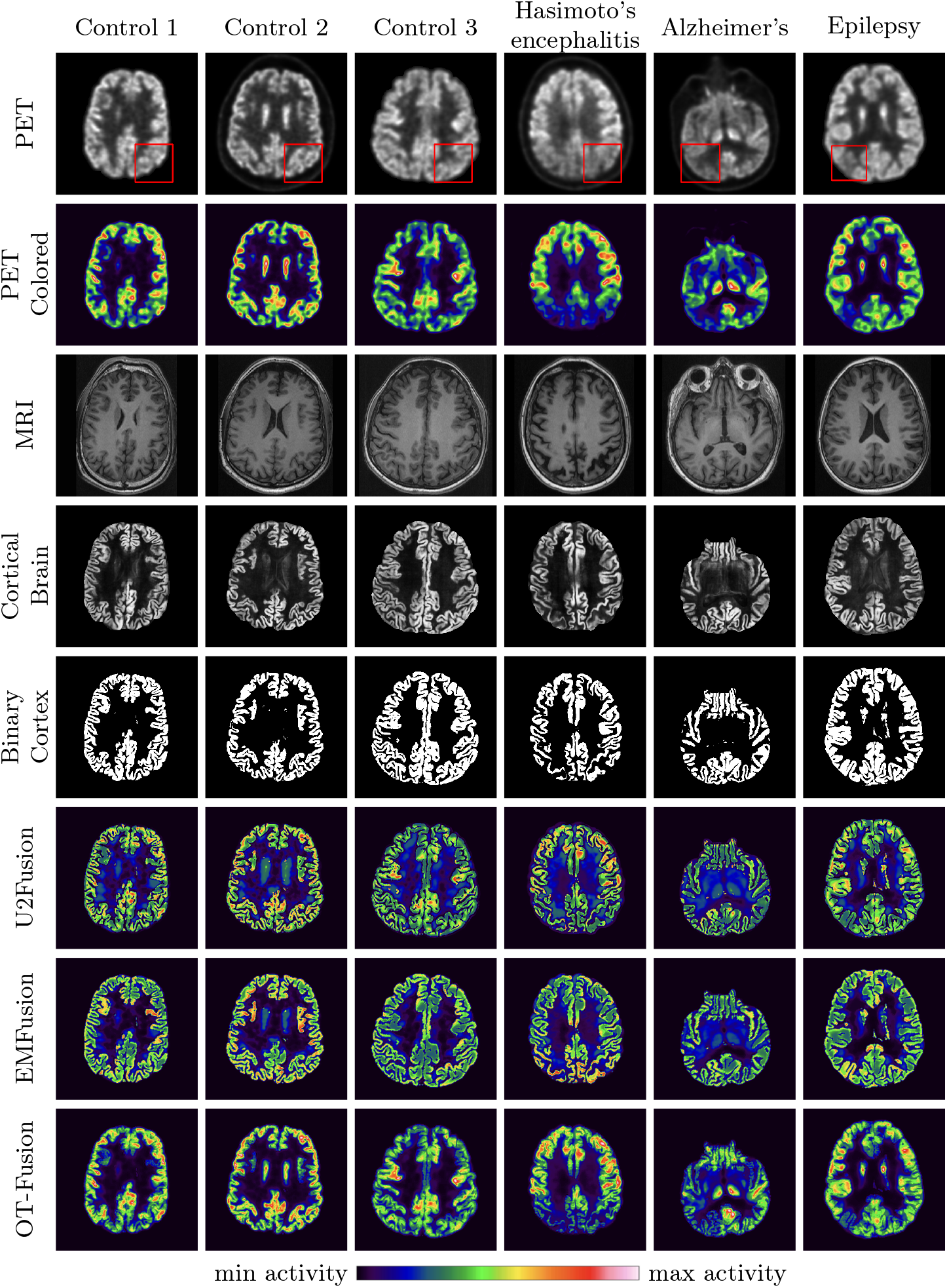
Comparison of image fusion methods across healthy (control) individuals (columns 1–3) and individuals with conditions (columns 4–6). For the first three columns red boxes are chosen randomly but for the last three columns red boxes indicate the affected regions due to the conditions that are zoomed in Figure 2. The color map used for representing the spectrum from minimum activity to maximum activity is shown below all images. The color scheme is called French-LUT.

**Fig. 2.**
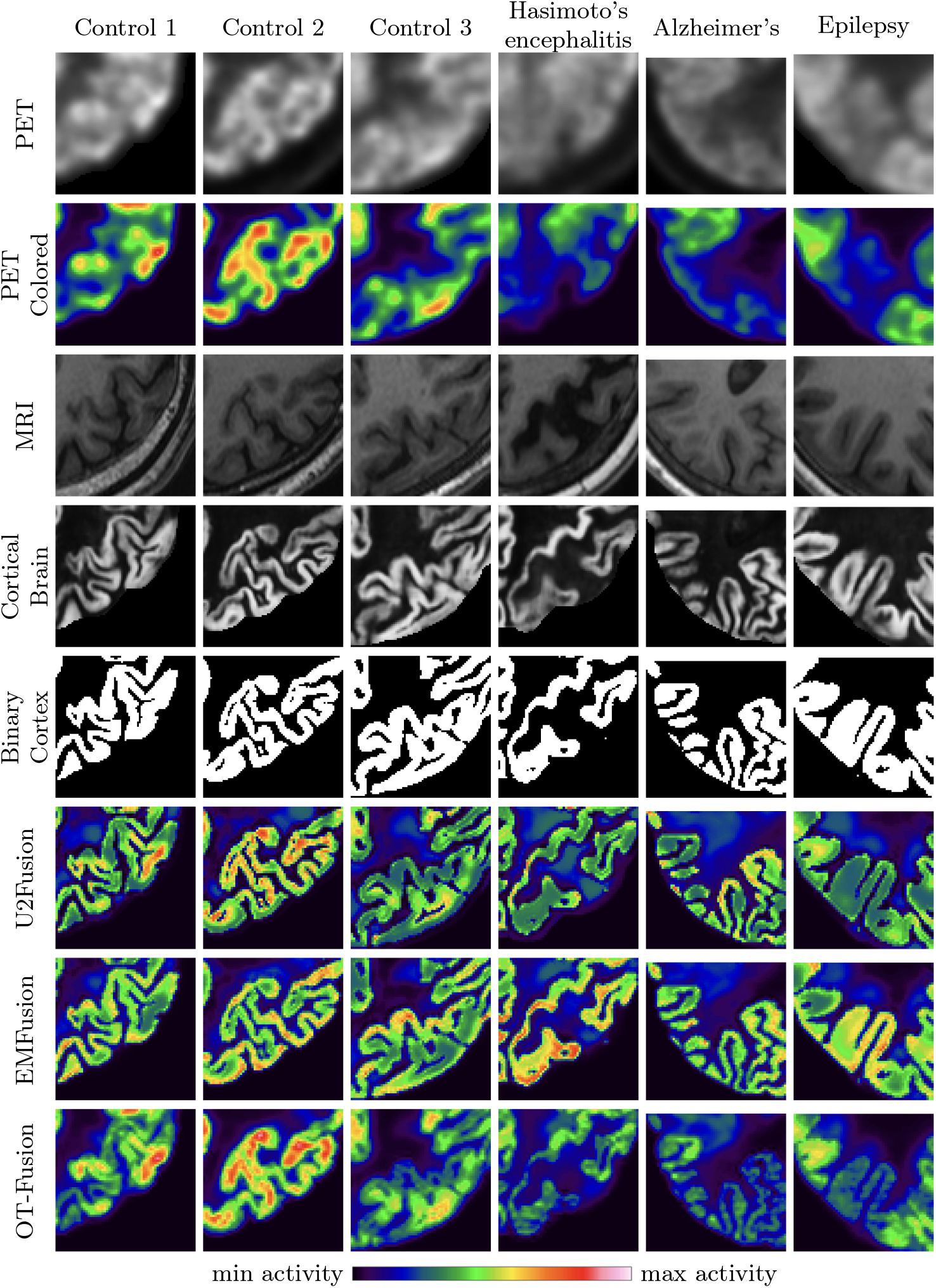
Zoomed-in versions of the images from Figure 1. These images correspond to the regions of interest marked with red boxes in the first row of that figure.

OT-Fusion accurately identifies hypometabolic cortical regions observed in original PET images, maintaining critical disease indicators. In contrast, the EMFusion method represents these regions as hypermetabolic, while the U2Fusion method depicts them as normometabolic or only mildly hypometabolic, potentially leading to diagnostic confusion. In the latter case (column 6), precise localization of metabolic abnormalities is crucial to the eventual surgical treatment of epilepsy. Accurate identification of these regions ensures that the epileptogenic zone is correctly targeted during resection, which is essential for achieving seizure freedom post-surgery.

## 5 Conclusion

For the first time, we formulate the problem of mapping PET activity (image intensity) distribution to predicted cortical regions in the MRI space as an optimal transport problem. Here, transport refers to the cost-effective transform of one probability distribution into another. OT-Fusion ensures that the transformed PET data aligns closely with the cortical region which we verify quantitatively and qualitatively. We show that OT-Fusion is an effective method that can be used by physicians when evaluating matched PET / MRI images.

## Footnotes

3 https://clinicaltrials.gov/study/NCT05355857

4 https://github.com/hanna-xu/U2Fusion

5 https://github.com/hanna-xu/EMFusion

